# The ubiquitin-conjugating enzyme UBE2D/eff maintains a youthful proteome and ensures protein quality control during aging

**DOI:** 10.1101/2023.12.12.571303

**Authors:** Liam C. Hunt, Kudzai Nyamkondiwa, Anna Stephan, Jianqin Jiao, Kanisha Kavdia, Vishwajeeth Pagala, Junmin Peng, Fabio Demontis

**Author notes:** Department of Biology, Rhodes College, 2000 North Pkwy, Memphis, TN 38112, USA.

## Abstract

Ubiquitin-conjugating enzymes (E2s) are key for regulating protein function and turnover via ubiquitination but it remains undetermined which E2s maintain proteostasis during aging. Here, we find that E2s have diverse roles in handling a model aggregation-prone protein (huntingtin-polyQ) in the *Drosophila* retina: while some E2s mediate aggregate assembly, UBE2D/effete (eff) and other E2s are required for huntingtin-polyQ degradation. UBE2D/eff is key for proteostasis also in skeletal muscle: eff protein levels decline with aging, and muscle-specific eff knockdown causes an accelerated buildup in insoluble poly-ubiquitinated proteins (which progressively accumulate with aging) and shortens lifespan. Transgenic expression of human UBE2D2, homologous to eff, partially rescues the lifespan and proteostasis deficits caused by muscle-specific eff^RNAi^ by re-establishing the physiological levels of eff^RNAi^-regulated proteins, which include several regulators of proteostasis. Interestingly, UBE2D/eff knockdown in young age reproduces part of the proteomic changes that normally occur in old muscles, suggesting that the decrease in UBE2D/eff protein levels that occurs with aging contributes to reshaping the composition of the muscle proteome. Altogether, these findings indicate that UBE2D/eff is a key E2 ubiquitin-conjugating enzyme that ensures protein quality control and helps maintain a youthful proteome composition during aging.

## INTRODUCTION

Protein degradation regulates many cellular functions, and its derangement is the underlying cause of many human diseases (*1-4*). Most proteins are regulated via degradation to ensure dynamic adjustments in their concentrations in response to cellular challenges and to maintain physiologic homeostasis. Moreover, damaged and misfolded proteins are degraded to avoid toxicity caused by their interaction with native proteins (*1–4*). Because of these important functions, protein degradation is tightly controlled. In most cell types, the ubiquitin/proteasome system (UPS) is responsible for most of the protein degradation in the nucleus and cytoplasm whereas the autophagy/lysosome system degrades cellular organelles, protein aggregates, and long-lived proteins (*1–6*). In addition, several proteases and peptidases cooperate with the proteasome and autophagy in degrading target proteins (*7–10*).

To route proteins for degradation, the UPS relies on ubiquitination, a post-translational modification that also regulates protein function and localization (*11, 12*). Through the concerted actions of a single ubiquitin-activating enzyme (E1), ∼35 ubiquitin-conjugating enzymes (E2s), and ∼620 E3 ubiquitin ligases, the ubiquitin-proteasome system (UPS) orchestrates the specific poly-ubiquitination of protein substrates in humans, which can lead to their degradation (*6, 13-16*). Depending on the ubiquitin lysine residue employed to build poly-ubiquitin chains, poly-ubiquitinated proteins can be preferentially degraded by the proteasome or by the autophagy-lysosome system (*11, 12, 17-21*). E2s have a pivotal role in the ubiquitination cascade and direct the recruitment of E3 ubiquitin ligases and target proteins (*21–24*).

In addition to regulating normal protein turnover, E2s can also ensure proteostasis. For example, the E2 enzyme UBE2B and its associated E3 ligase UBR4 have been found to regulate muscle protein quality during aging in *Drosophila* and mice (*25, 26*). The capacity of E2s to maintain proteostasis may derive from their capacity to guide the ubiquitination and proteasome-mediated degradation of misfolded and aggregation-prone proteins (*27, 28*) and to trigger autophagy (*29, 30*). Moreover, because some E2s are physically associated with the proteasome, it has been proposed that they may directly regulate its proteolytic activity (*31–33*). For example, the E2 enzymes Ubc1/2/4/5 interact with the proteasome and this association further increases with heat stress in yeast (*31*), a condition where proteasome function is modulated by ubiquitination of proteasome components, such as Rpn13, mediated by the proteasome-associated E3 UBE3C (*34*). While E2s may regulate proteostasis via several mechanisms, it remains largely undetermined which E2s are crucial for maintaining protein quality control during aging.

Here, we have utilized a model aggregation-prone protein (GFP-tagged pathogenic huntingtin, Htt-polyQ-GFP) expressed in the *Drosophila* retina to test the impact of E2 ubiquitin-conjugating enzymes on proteostasis. In addition, we have similarly tested a set of E3 ubiquitin ligases and deubiquitinating enzymes (DUBs) that were identified by mapping the E2 interactome in human cells (*35*). These analyses reveal diverse functions of E2s and associated enzymes in proteostasis. Contrary to the expectation that E2s may generally promote the proteasomal degradation of Htt-polyQ and hence reduce Htt-polyQ levels, we find that the knockdown of several E2s has the opposite effect, i.e. it decreases Htt-polyQ aggregates.

Knockdown of the E2 enzyme effete (eff), homologous to human UBE2D1/2/3/4 ubiquitin-conjugating enzymes, is one of the relatively few RNAi interventions that increase the amount of Htt-polyQ aggregates, and we find that this is due to an impediment in Htt-polyQ protein degradation. Consistent with a prevalent role of UBE2D/eff in proteostasis across aging tissues in *Drosophila*, we find that UBE2D/eff protein levels significantly decline during aging in skeletal muscle, and that muscle-specific eff knockdown increases the age-associated accumulation of insoluble poly-ubiquitinated proteins and reduces lifespan. Interestingly, the proteostasis deficits caused by UBE2D/eff knockdown are partially rescued by transgenic expression of one of its human homologs, UBE2D2, suggesting that UBE2D ubiquitin-conjugating enzymes have evolutionary conserved roles in maintaining protein quality control during aging. Altogether, our study identifies diverse roles for E2 ubiquitin-conjugating enzymes in handling aggregation-prone proteins and managing proteostasis, and that UBE2D/eff is key for ensuring protein quality control during aging.

## RESULTS

### Diverse roles of E2 ubiquitin-conjugating enzymes in the disposal of pathogenic huntingtin

Huntington disease (HD) is caused by pathogenic huntingtin with polyQ tract expansion (Htt-polyQ), a toxic, aggregation-prone protein that induces neurodegeneration (*36–38*). Previous studies have shown that ubiquitination is a conserved regulator of polyglutamine aggregation and that E2 loss can both increase and decrease the size and number of Htt-polyQ aggregates (*25, 26, 39, 40*). However, as for other analyses of E2 function, a complete understanding of the role of E2s in polyglutamine aggregation is missing.

The fruit fly *Drosophila melanogaster* is an ideal model for studying the mechanisms of Htt-polyQ protein degradation. GFP-tagged Htt-polyQ72 can be expressed in the fly retina (*41*) and fluorescent Htt-polyQ-GFP aggregates can be visualized from intact animals over time (*42, 43*). In particular, the fly retina is a convenient tissue to examine Htt-polyQ because this tissue is dispensable for fly survival. With this model, it was previously found that there is an age-dependent increase in the amount of Htt-polyQ72 protein aggregates, which reflects the overall age dependency of HD phenotypes also observed in humans (*36, 37*).

To test whether E2s regulate Htt-polyQ-GFP aggregation during aging, multiple RNAi lines were utilized to target each of the 21 *Drosophila* E2s. For these studies, transgenic RNAi was driven in the *Drosophila* retina with the UAS/Gal4 system and GMR-Gal4, concomitantly with Htt-polyQ-GFP (Fig. 1A). After 30 days at 29°C, the total area of Htt-Q72-GFP fluorescent aggregates was scored with CellProfiler and compared to control RNAi interventions (Fig. 1A-C and Supplementary Table 1). Overall, there was a trend towards decreased aggregate area upon knockdown of several E2s, including the *Drosophila* homologs of UBE2F/M, UBE2I, UBE2QL1, and UBE2Z, suggesting that these E2s may be required for ubiquitination-dependent aggregation of Htt-polyQ-GFP (Fig. 1A-B). Analyses of deubiquitinating enzymes (DUBs) and E3 ubiquitin ligases identified from mapping the E2 interactome in human cells (*35*) indicate that RNAi for these interactors yields overall similar phenotypes as those found by knocking down the interacting E2s (Fig. 1A-C). For example, RNAi for E3s and DUBs that interact with Ubc10 (UBE2L3) and CG7656 (UBE2R1/2) reduce the area of Htt-polyQ-GFP aggregates, as found for knockdown of these E2s (Fig. 1C). The decreased polyglutamine aggregation that is observed upon E2 RNAi may indicate that E2-mediated ubiquitination is required for Htt aggregate formation. Consistent with this model, it was previously reported that K48 linkage-specific ubiquitination drives Htt degradation by the proteasome whereas K63 ubiquitination promotes Htt aggregation (*44*). Our comprehensive analysis now highlights the differential effects of distinct E2s on Htt disposal and suggests that this may occur because of the specificity in the E3s and DUBs interacting with each E2.

**Fig. 1.**
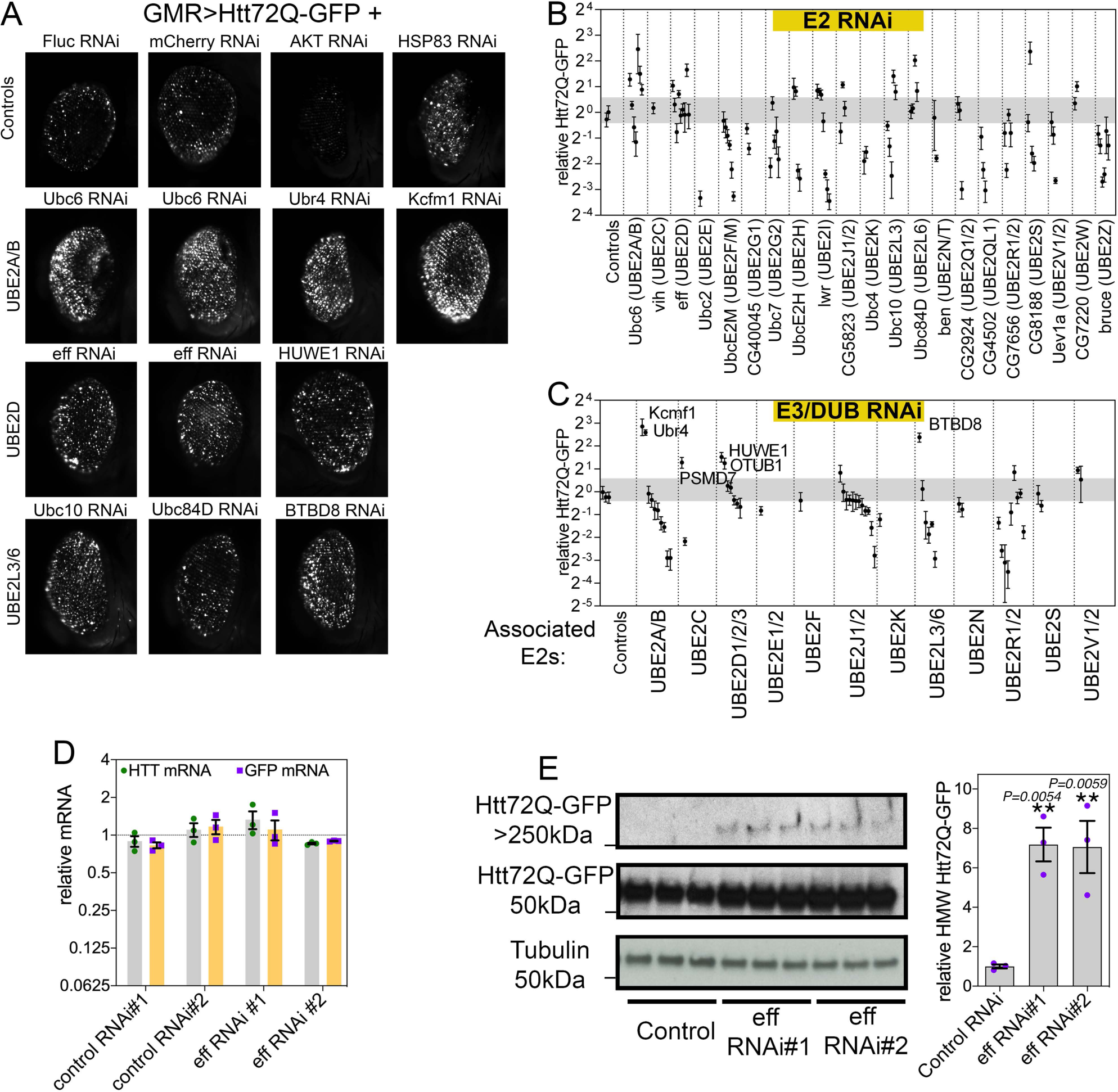
RNAi for E2 ubiquitin-conjugating enzymes and associated E3s and DUBs modulates pathogenic huntingtin-polyQ aggregates in the *Drosophila* retina. (**A**) GFP-tagged huntingtin-polyQ (Htt-polyQ72-GFP) driven with *GMR-Gal4* leads to GFP-fluorescent Htt protein aggregates in the retina at 30 days of age. Compared to negative controls (mCherry^RNAi^ and luciferase^RNAi^), RNAi for E2 ubiquitin-conjugating enzymes, associated deubiquitinating enzymes (DUBs), and E3 ubiquitin ligases (E3s) modulates the amount of Htt-polyQ72-GFP aggregates. Specifically, RNAi for Ubc6 (homologous of UBE2A/B) and for its associated E3 ubiquitin ligases Ubr4 and Kcmf1 increases the amount of Htt-polyQ72-GFP protein aggregates. Similar increases are also seen with knockdown of eff/UBE2D and the associated E3 HUWE1, and with RNAi for Ubc10/UBE2L3, Ubc84D/UBE2L6, and for the associated E3 enzyme BTBD8. Positive controls include Akt RNAi, which reduces protein aggregates, and Hsp83 RNAi, which increases them. (**B-C**) Quantitation of the total area of Htt-polyQ72-GFP aggregates modulated by RNAi for E2s (B) and for associated E3s and DUBs (C). Relative fold changes compared to control RNAi interventions are shown; n=5 (biological replicates), SD. Each data point in the graph represents a single RNAi targeting the corresponding gene. For each RNAi, the mean of 5 biological replicates is shown, with each biological replicate representing a single eye (each from a distinct animal). **(D)** qRT-PCR indicates that there are no changes in *GFP* and *Htt* mRNA levels upon eff knockdown compared to control RNAi, indicating that eff RNAi does not modulate the amount of GFP-tagged huntingtin-polyQ aggregates via changes in the expression of *Htt-polyQ72-GFP* transgenes; n=3 (biological replicates) and SEM. **(E)** Levels of Htt-polyQ72-GFP aggregates detected by western blot with anti-GFP antibodies identify Htt-polyQ72-GFP monomers (∼50 kDa) and high-molecular-weight (HMW) assemblies of Htt-polyQ72-GFP in the stacking gel (>250 kDa). Knockdown of eff/UBE2D increases the levels of HMW Htt-polyQ72-GFP; n=3 (biological replicates), SEM, and *p*-values (one-way ANOVA) are indicated, with \*\**p*<0.01, compared to mcherry^RNAi^.

### UBE2D/eff knockdown increases aggregates of pathogenic huntingtin in the *Drosophila* retina

In addition to RNAi lines that decrease the amount of Htt aggregates, we found that RNAi-mediated knockdown of other E2s causes an increase in Htt aggregates (Fig. 1A-C). Specifically, RNAi for Ubc6 (UBE2A/B), eff (UBE2D), Ubc10 (UBE2L3), and Ubc84D (UBE2L6) increased the total area of protein aggregates. Likewise, RNAi for E3/DUBs that physically associate with these E2s (such as Ubr4 and Kcfm1, associated with UBE2A/B; and HUWE1, associated with UBE2D) led to higher levels of polyglutamine aggregates. A possible interpretation of these results is that these E2s are necessary for the degradation of aggregation-prone Htt and, consequently, their loss leads to the accumulation of detergent-insoluble polyglutamine aggregates. To test this hypothesis, we further characterized the molecular mechanisms by which eff/UBE2D regulates Htt proteostasis. We first measured by qRT-PCR the levels of Htt-polyQ-GFP and found that there are no changes in its expression upon eff RNAi compared to controls (Fig. 1D), indicating that increased levels of Htt-polyQ-GFP aggregates observed upon eff RNAi do not arise from increased transgenic expression of Htt-polyQ-GFP. Next, we utilized western blot to analyze the levels of Htt-polyQ-GFP monomers (∼50 kDa) and high-molecular-weight (HMW) Htt-polyQ-GFP aggregates that accumulate in the stacking gel (>250 kDa). Remarkably, 2 different RNAi lines targeting eff/UBE2D increase HMW Htt-polyQ-GFP levels (Fig. 1E), suggesting that eff/UBE2D-mediated ubiquitination may be required for Htt degradation. In summary, these findings indicate an extensive role of E2 ubiquitin-conjugating enzymes in modulating Htt-Q72-GFP aggregation during aging in the *Drosophila* retina.

### Retinal degeneration caused by UBE2D/eff knockdown is rescued by human UBE2D2/4

Compared to controls, we find that RNAi for Ubc6 and for eff induces retinal degeneration, as indicated by the “rough eye” appearance (indicative of photoreceptor death and derangement of inter-ommatidial bristles (*45, 46*)), the increase in the eye area occupied by necrotic patches, and/or the loss of pigmentation (Fig. 2A). On this basis, we next examined whether concomitant transgenic expression of the human homologs of these *Drosophila* E2s rescues retinal degeneration. On this basis, we expressed human UBE2B (homologous to Ubc6, DIOPT homology score=13) and human UBE2D2 and UBE2D4 (homologous to eff, DIOPT scores of 13 and 10, respectively) and compared their capacity to rescue retinal degeneration compared to mock mcherry overexpression (Fig. 2A). As expected based on the sequence homology, hUBE2B (but not mcherry, hUBE2D2, and hUBE2D4) rescued retinal degeneration (i.e. necrotic patches and depigmentation) induced by Ubc6 RNAi. Similarly, hUBE2D2 and to a lower extent hUBE2D4 rescued depigmentation caused by eff knockdown. Altogether, these findings indicate that Ubc6 and eff loss can be rescued by transgenic expression of their respective human homologs UBE2B and UBE2D2/4 (Fig. 2A).

**Fig. 2.**
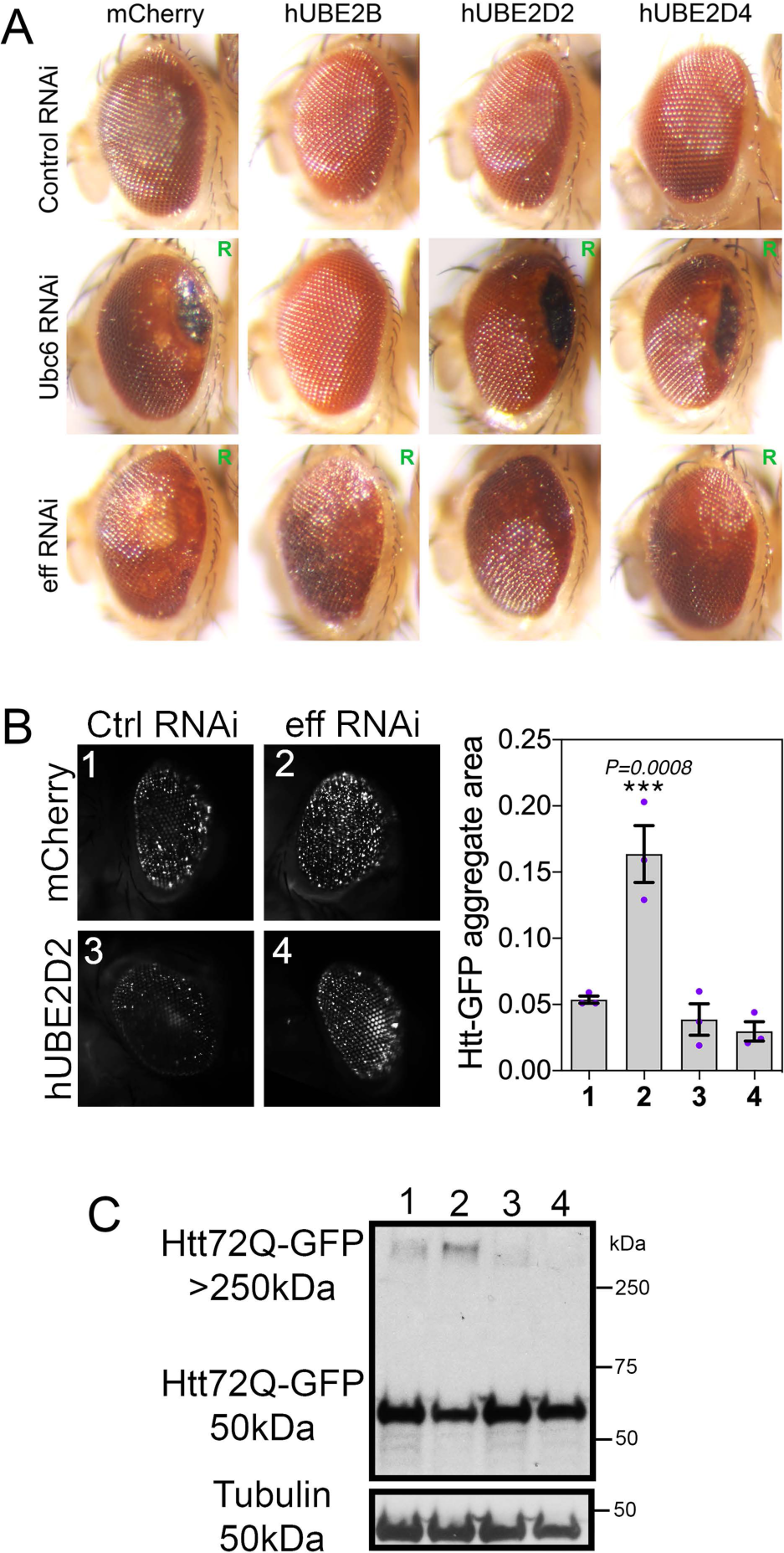
Retinal degeneration induced by knockdown of Ubc6 and eff is rescued by expression of the respective human homologs UBE2B and UBE2D2/4. (**A**) Compared to control RNAi, knockdown of Ubc6 and eff causes retinal degeneration, as indicated by the “rough eye” phenotype (R), by black necrotic patches, and/or by areas with depigmentation. Human UBE2B rescues degeneration induced by RNAi for its *Drosophila* homolog Ubc6 but not induced by RNAi for the unrelated E2 enzyme eff. Likewise, human UBE2D2 and UBE2D4 rescue retinal degeneration induced by knockdown of their *Drosophila* homolog eff. (**B**) Htt-polyQ72-GFP aggregates increase in *Drosophila* retinas with eff RNAi and this is rescued by human UBE2D2 but not by mcherry; n=3 (biological replicates), SEM, and *p*-values (one-way ANOVA) are indicated, with \*\*\**p*<0.001. (**C**) Similarly, high-molecular-weight assemblies of Htt-polyQ72-GFP detected in the stacking gel increase with eff RNAi compared to control RNAi but this is rescued by hUBE2D2 versus control mcherry expression.

On this basis, we next examined whether hUBE2D can rescue the defects in Htt-polyQ proteostasis caused by eff RNAi. Specifically, we compared eff RNAi versus control RNAi, with the concomitant expression of either mcherry or hUBE2D2. Image analysis of Htt-poly-GFP aggregates indicates that hUBE2D2 impedes the accumulation of Htt-polyQ-GFP aggregates which are otherwise increased by eff RNAi (Fig. 2B). Similar results were also obtained by western blot: there was lower accumulation of HMW Htt-polyQ-GFP in eff^RNAi^+hUBE2D2 versus eff^RNAi^+mcherry and the other controls (Fig. 2C). In summary, these findings indicate that eff/UBE2D has an evolutionary-conserved function in maintaining proteostasis in a Huntington’s disease model.

### UBE2D/eff protein levels decline with aging, and UBE2D/eff knockdown impairs skeletal muscle proteostasis as observed during aging

The comprehensive analysis reported above has revealed a range of diverse roles for E2s in modulating polyglutamine aggregates in *Drosophila*. UBE2D/eff appears to be one of the E2s with the most striking effects in preserving proteostasis, i.e. it is required for preventing the accumulation of HMW Htt-polyQ-GP (Fig. 1-2). On this basis, we further probed the role of UBE2D/eff in protein quality control. To this purpose, we examined skeletal muscle aging, which is characterized by a progressive decline in proteostasis and by the age-related accumulation of detergent-insoluble poly-ubiquitin protein aggregates (*25, 47-53*).

Deep-coverage tandem mass tag (TMT) mass spectrometry (*54, 55*) of skeletal muscle from young and old control w^1118^ flies (1 and 8 weeks old) detected 5,971 proteins, including 10 of the 21 *Drosophila* E2 ubiquitin-conjugating enzymes (Fig. 3A and Supplementary Table 2). Interestingly, UBE2D/eff displayed the strongest age-dependent downregulation (Fig. 3B), suggesting that a reduction in UBE2D/eff protein levels may contribute to the progressive loss of protein quality control with aging.

**Fig. 3.**
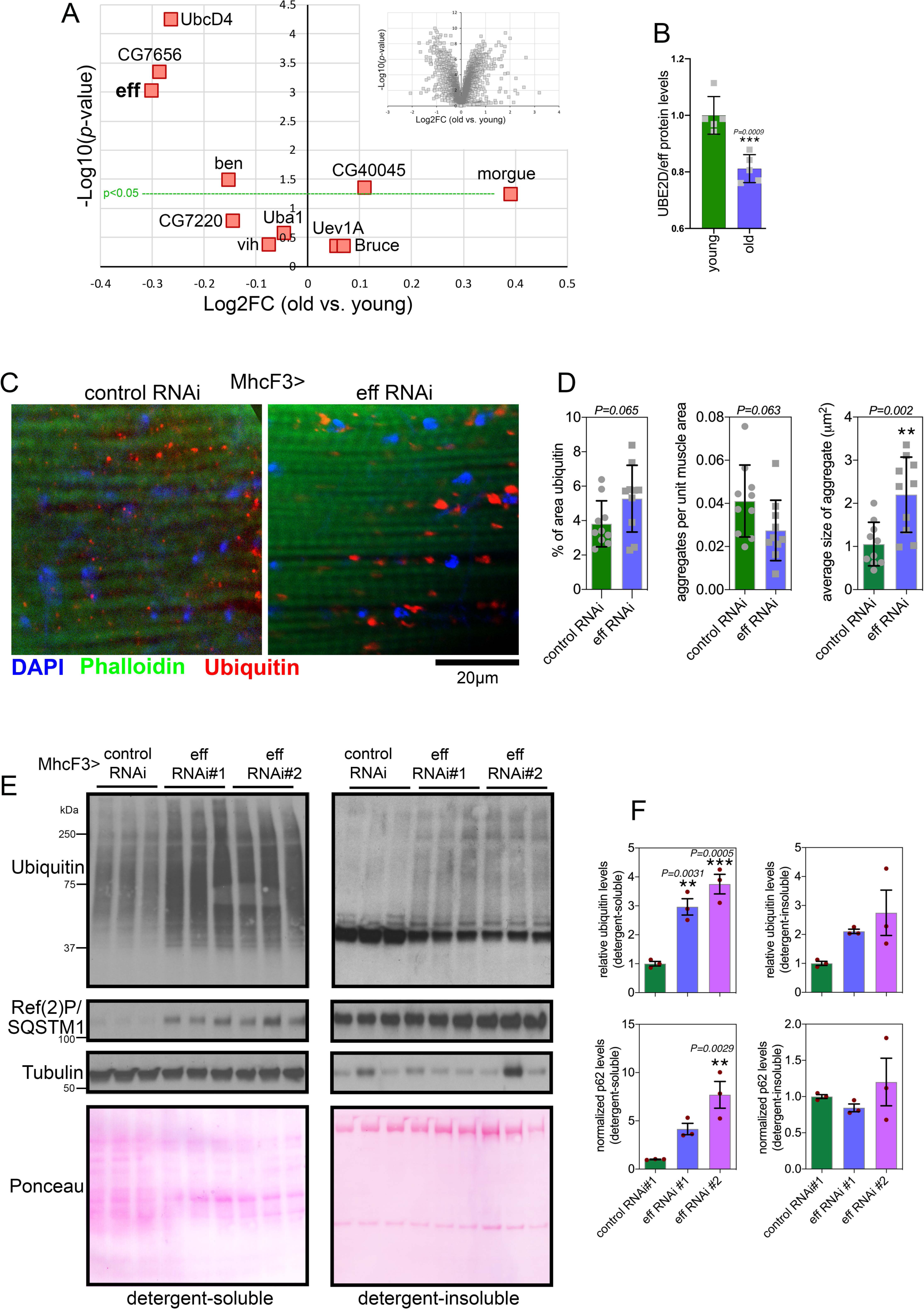
RNAi for the E2 enzyme eff/UBE2D impairs muscle protein quality control during aging. **(A)** TMT mass spectrometry of *Drosophila* skeletal muscle from old versus young w^1118^ flies at 8 vs. 1 week old; n=5 biological replicates. The *x* axis reports the Log2FC whereas the *y* axis reports the –log10(*p*-value). The E1 enzyme UBA1 and the E2 enzymes that are detected are indicated, including UBE2D/eff; n=5 (biological replicates). **(B)** The protein levels of UBE2D/eff significantly decline with aging (8 versus 1 week-old) in skeletal muscle; n=5 (biological replicates), SD, and *p*-values (Student’s *t*-test). (**C-D**) Immunostaining of *Drosophila* skeletal muscle at 3 weeks of age indicates that eff/UBE2D RNAi impairs protein quality control, as indicated by the higher age-related accumulation of aggregates of poly-ubiquitinated proteins, compared to control RNAi. The scale bar is 20 μm. In (C), n=10 (biological replicates), SD, and *p*-values (Student’s *t*-test). (**E-F**) Western blot analysis of detergent-soluble and insoluble fractions from *Drosophila* skeletal muscle indicates that eff RNAi impedes proteostasis, as indicated by higher levels of detergent-soluble and insoluble poly-ubiquitinated proteins compared to control RNAi (E). A similar increase is also found for the detergent-soluble levels of Ref(2)P/p62 (E). n=3 (biological replicates), SEM, \*\**p*<0.01, ns=not significant (one-way ANOVA).

To test this hypothesis, we reduced UBE2D/eff levels from young age by driving UBE2D/eff RNAi with MhcF3-Gal4 (*25*) in skeletal muscle, and compared this intervention to a control mcherry RNAi. Immunostaining and confocal microscopy of skeletal muscle of flies at 3 weeks of age indicate that UBE2D/eff RNAi significantly increases the size of poly-ubiquitin protein aggregates and that there is an overall trend towards a higher total area of aggregates, compared to control RNAi (Fig. 3C-D). Altogether, these results indicate that anticipating the age-related decline in eff/UBE2D protein levels deranges proteostasis.

We next examined muscle proteostasis via the analysis of detergent-soluble and insoluble fractions (*56*). Knockdown of UBE2D/eff with 2 distinct RNAi lines increased the levels of poly-ubiquitinated proteins found in both the soluble and insoluble fractions of skeletal muscle of 3-weeks-old flies. There was also an increase in Ref(2)P/p62 levels that occurred in parallel with the accumulation of poly-ubiquitinated proteins, although this occurred primarily in the detergent-soluble fraction (Fig. 3E-F). Altogether, these findings indicate that UBE2D/eff is required to ensure protein quality control in skeletal muscle.

### Defects in proteostasis and lifespan caused by muscle-specific UBE2D/eff knockdown are partially rescued by transgenic expression of human UBE2D2

We have found that human UBE2D2 can rescue retinal degeneration induced by eff RNAi in the context of polyglutamine disease, and that this results from the degradation of HMW Htt-polyQ (Fig. 2). On this basis, we next examined whether hUBE2D2 can also rescue defects in muscle proteostasis due to eff knockdown. To this purpose we examined detergent-soluble and insoluble fractions from muscles of flies with eff^RNAi^+hUBE2D2 versus eff^RNAi^+mcherry as well as control^RNAi^+hUBE2D2 versus control^RNAi^+mcherry (Fig. 4A-B). Western blot analyses with anti-ubiquitin and anti-p62/Ref(2)P antibodies revealed that eff^RNAi^+hUBE2D2 reduced the detergent-soluble levels of ubiquitinated proteins and p62/Ref(2)P compared to eff^RNAi^+mcherry and other controls, whereas there was no rescue of detergent-insoluble levels (Fig. 4A-B).

**Fig. 4.**
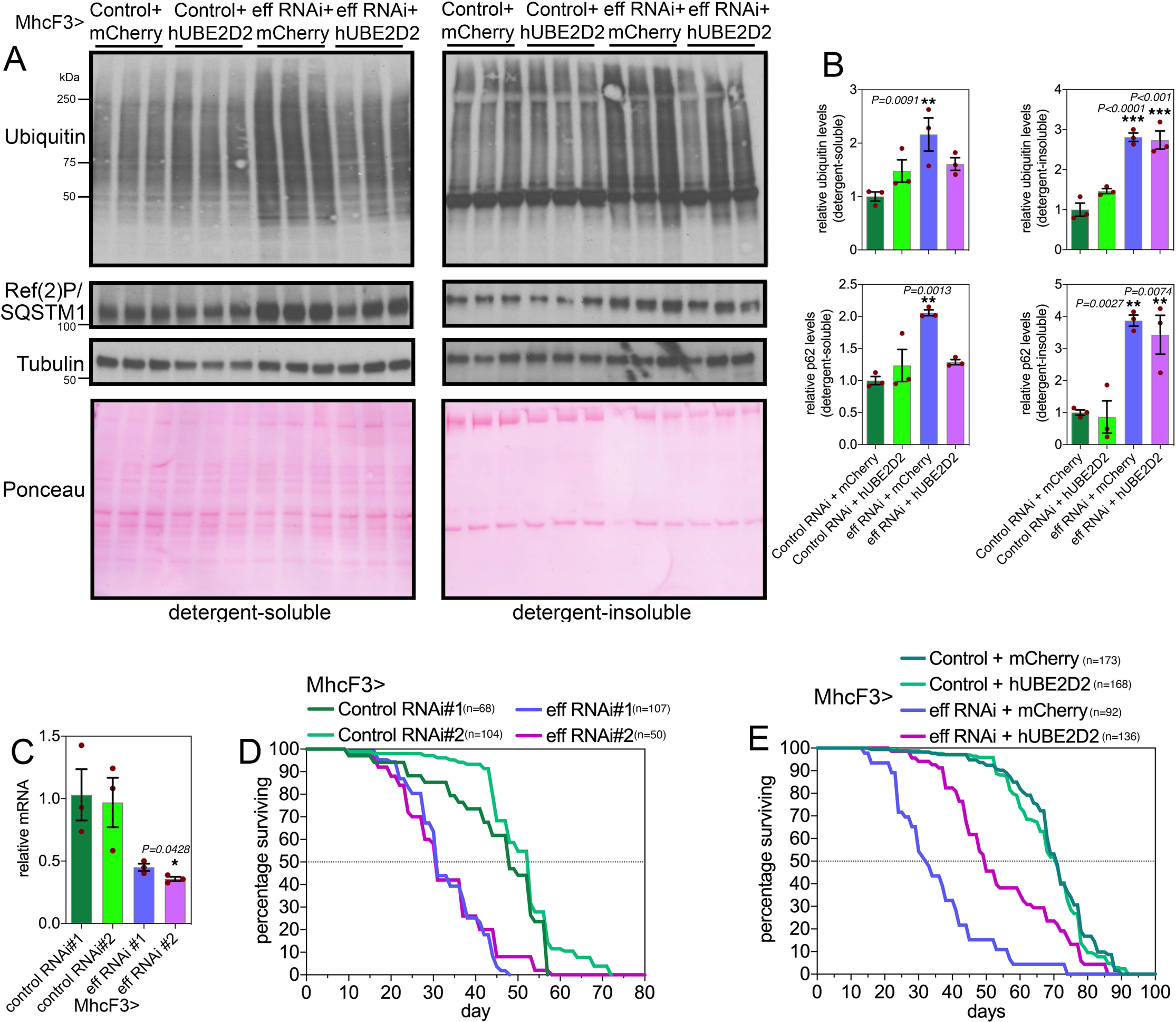
Defects in proteostasis and lifespan caused by muscle-specific eff/UBE2D RNAi are partially rescued by the expression of human UBE2D2. (**A-B**) Western blot analysis of detergent-soluble and insoluble fractions from *Drosophila* skeletal muscle indicates that defects in proteostasis due to eff RNAi can be partially rescued by its human homolog UBE2D2, as indicated by the normalization of the detergent-soluble levels of poly-ubiquitinated proteins and Ref(2)P/p62. However, hUBE2D2 does not impact their detergent-insoluble levels, indicating that hUBE2D only partially rescues defects induced by eff/UBE2D RNAi. In (B), n=3 (biological replicates), SEM, \*\**p*<0.01 (one-way ANOVA). (**C**) Muscle-specific eff/UBE2D RNAi reduces eff mRNA levels; n=3 (biological replicates), SEM, \*\**p*<0.01 (one-way ANOVA). (**D**) Two distinct RNAi lines targeting eff/UBE2D reduce lifespan compared to control RNAi (*p*<0.001, log-rank test). (**E**) The decline in organismal survival due to eff/UBE2D RNAi in muscle is partially rescued by transgenic hUBE2D expression compared to control mcherry overexpression (*p*<0.001, log-rank test).

Because muscle proteostasis has been found to regulate organismal aging (*25, 47, 49, 57, 58*), we next determined the impact of muscle-specific UBE2D/eff loss on lifespan. Knockdown of eff with 2 distinct RNAi lines (Fig. 4C) shortened lifespan (Fig. 4D) and this was partially rescued by concomitant expression of hUBE2D but not by control mcherry (Fig. 4E). Altogether, these studies identify a key role for UBE2D/eff in proteostasis during skeletal muscle aging in *Drosophila*, and highlight the evolutionary conservation of UBE2D function.

### Deep-coverage proteomics reveals protein substrates that are modulated by eff/UBE2D

To determine the proteins that are modulated by UBE2D/eff RNAi and the partial rescue of proteostasis defects by hUBE2D, TMT-based proteomics (*54, 55*) was utilized to determine the protein changes modulated by eff/UBE2D RNAi versus control RNAi, and whether these are rescued by hUBE2D2 compared to control mcherry expression (*35*). Analysis of eff-induced protein changes revealed several categories that are enriched among the significantly regulated proteins (Fig. 5A-D and Supplementary Table 3; (*35*)). Upregulated proteins included proteasome components, chaperones, deubiquitinases, ubiquitin ligases, and extracellular proteins whereas downregulated proteins were enriched for peptidases, lipases, and secreted proteins (Fig. 5E).

**Fig. 5.**
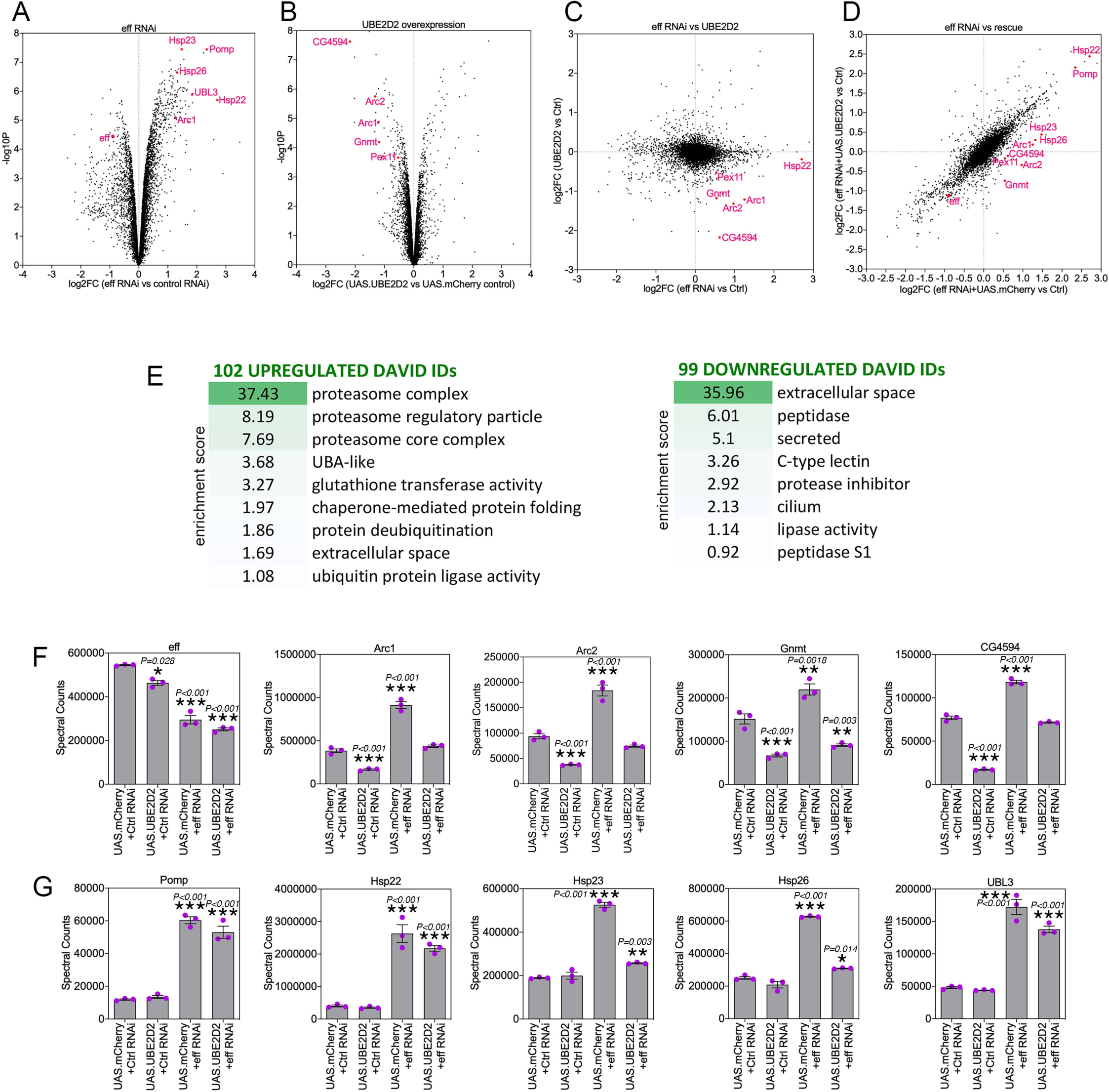
Deep-coverage TMT mass spectrometry identifies the proteins modulated by UBE2D/eff knockdown in *Drosophila* skeletal muscle. (**A-B**) TMT mass spectrometry of *Drosophila* skeletal muscle with eff/UBE2D knockdown versus control RNAi (A) and overexpression of human UBE2D2 versus control mcherry (B). The *x* axis reports the Log2FC whereas the *y* axis reports the –log10(*p*-value). Examples of regulated proteins are shown in red. (**C**) Cross-comparison of the Log2FC induced by eff RNAi versus control RNAi (*x* axis) with the Log2FC of hUBE2D2 versus control mcherry (*y* axis). (**D**) Cross-comparison of the Log2FC induced by eff RNAi + mcherry versus control (*x* axis) with the Log2FC of eff RNAi + hUBE2D2 versus control (*y* axis) indicates that some of the protein changes induced by eff RNAi are rescued by hUBE2D2. (**E**) GO term analysis of protein categories that are enriched among upregulated and downregulated proteins in response to eff/UBE2D2 RNAi. The enrichment scores and number of regulated DAVID IDs are shown. (**F-G**) Examples of regulated proteins modulated by eff RNAi and hUBE2D-mediated rescue include protein changes that may drive derangement of proteostasis, such as Arc1/2 upregulation (F) as well as protein changes that are protective and likely compensatory, such as proteasome components and chaperones (G); n=3 (biological replicates), SEM, \**p*<0.05, \*\**p*<0.01, \*\*\**p*<0.001 (one-way ANOVA).

Curated analyses revealed that, as expected, eff protein levels declined in response to eff RNAi compared to control RNAi (Fig. 5F). The levels of several other proteins were increased upon eff RNAi but reduced by concomitant rescue with hUBE2D2 (Fig. 5F). These proteins include Arc1 and Arc2 (activity-regulated cytoskeleton-associated protein 1 and 2), which regulate starvation-induced locomotion, the neuromuscular junction, and metabolism (*59–61*); the glycine N-methyltransferase Gnmt enzyme, which generates sarcosine and controls the amount of the methyl donor S-adenosylmethionine (*62, 63*); and CG4594, which encodes for an enzyme involved in fatty acid beta oxidation. In addition to rescuing protein changes induced by eff RNAi, overexpression of hUBE2D2 by itself reduced the levels of Arc1, Arc2, Gnmt, and CG4594. Altogether, these proteomics analyses identify protein targets that are modulated by eff/UBE2D in skeletal muscle (Fig. 5F). Improper eff/UBE2D-mediated turnover of these proteins and their consequent accumulation may contribute to derange muscle protein quality control and in turn to reduced survival. In agreement with this hypothesis, it was previously found that elevation of Arc1 protein levels in Alzheimer’s disease models causes cytotoxicity and contributes to neuronal death in *Drosophila* (*64*).

Conversely, there were other proteins that were induced in muscles with eff/UBE2D knockdown and that may represent a protective response that mitigates the derangement of proteostasis due to eff/UBE2D knockdown. These included Pomp, involved in proteasome assembly (*65, 66*), the chaperones Hsp22, Hsp23, and Hsp26, and the ubiquitin-like protein UBL3 (Fig. 5G).

In summary, these studies indicate that eff/UBE2D knockdown induces several proteomics changes, some of which are rescued by hUBE2D2. While some of these changes may be cytotoxic (e.g. increased Arc1 levels, (*64*)), others may conversely limit the derangement of protein quality control and/or improve survival.

### RNAi for UBE2D/eff causes proteomic changes associated with aging in *Drosophila* skeletal muscle

We have found that UBE2D/eff protein levels decline during aging in skeletal muscle (Fig. 3A-B) and that the experimental induction of UBE2D/eff knockdown in young age induces a premature decline in proteostasis (Fig. 3-4), which otherwise is normally lost only in old age (*47, 48*). Lastly, we have found that RNAi for UBE2D/eff induces many proteomic changes (Fig. 5), consistent with its role in ubiquitin-dependent protein regulation. On this basis, we next examined how the proteomic changes induced by aging correlate with those induced by eff RNAi in young age. To this purpose, we cross-compared the TMT mass spectrometry datasets obtained from *Drosophila* skeletal muscle with eff RNAi (normalized by control mcherry RNAi) and those obtained from control muscles of old versus young flies. Cross-comparison of the significantly regulated genes revealed that 700 out of 1,002 proteins (∼70%) are consistently regulated by eff RNAi and aging, whereas the remaining 302 proteins are discordantly regulated (Fig. 6A-D and Supplementary Table 4).

**Fig. 6.**
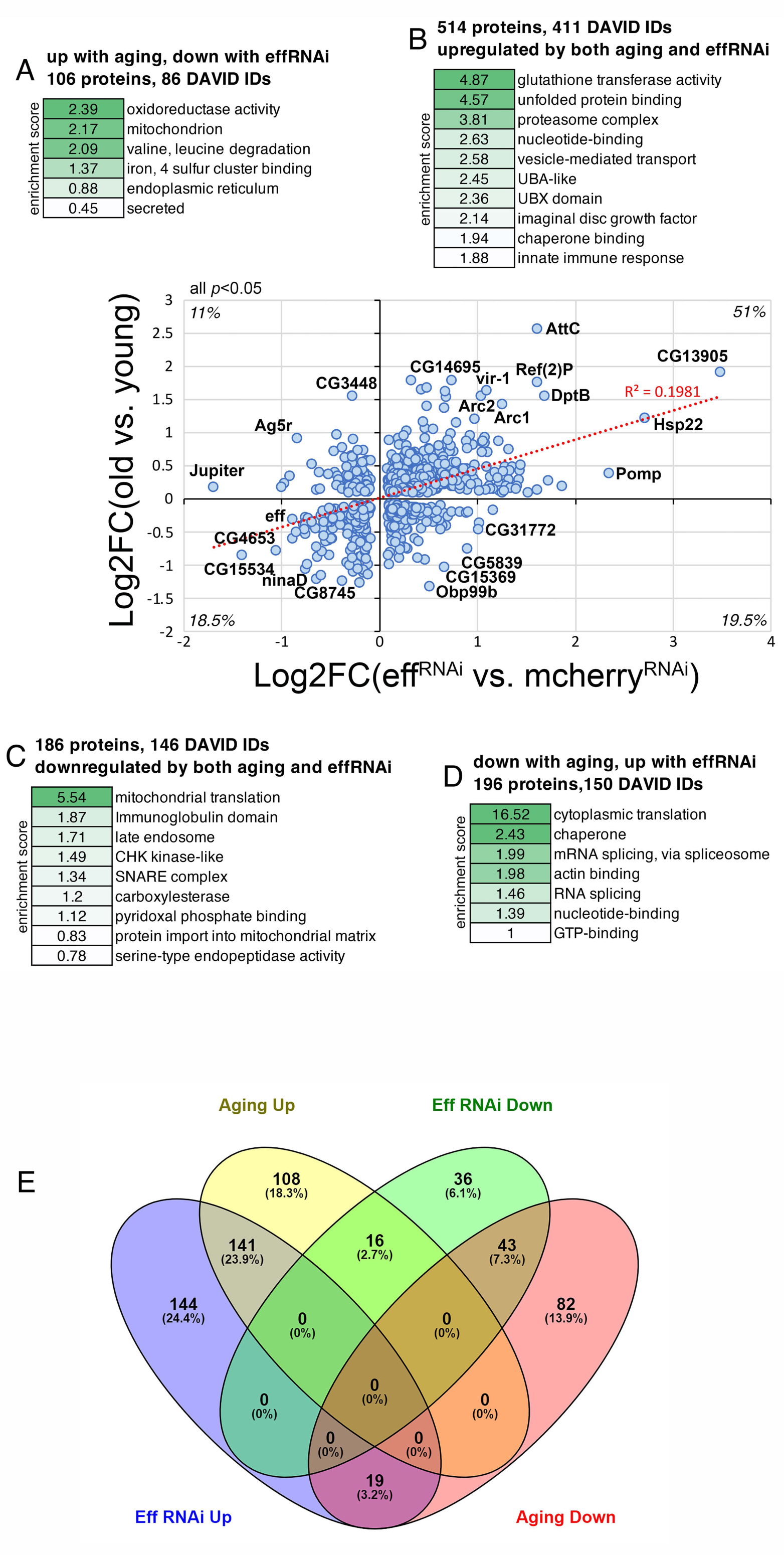
Knockdown of the ubiquitin-conjugating enzyme UBE2D/eff drives proteomic changes associated with aging in *Drosophila* skeletal muscle. (A-D) Cross-comparison of TMT mass spectrometry data from *Drosophila* skeletal muscle identifies substantial overlap in the proteomic changes induced by UBE2D/eff knockdown in young age compared to the changes that are induced by aging. The *x* axis displays the significant (*p*<0.05) changes (log_2_FC) induced in skeletal muscle by eff RNAi (*Mhc>eff^RNAi^*) compared to control mcherry RNAi (*Mhc>mcherry^RNAi^*) at 2 weeks of age (n=3 biological replicates/group). The *y* axis reports the significant (*p*<0.05) changes induced by aging in the skeletal muscle of control flies (*w^1118^*) when comparing 8 weeks (old) versus 1 week (young), with n=5 biological replicates/group. Among the proteins that are significantly regulated (*p*<0.05) by both UBE2D/eff knockdown and aging, ∼70% are consistently regulated, i.e. either upregulated (B, 51%) or downregulated (A, 18.5%) by both, whereas the remaining ∼30% is regulated oppositely by UBE2D/eff RNAi versus aging (A, D). The protein categories that are over-represented in each group are indicated in (A-D) alongside the enrichment score. Representative proteins that are significantly regulated by UBE2D/eff RNAi in a consistent or discordant manner are shown in the graph and include Arc1 and Arc2. **(E)** Venn diagrams representing the overlap in the regulation of protein levels by aging and eff^RNAi^. These graphs were obtained from the list of significantly regulated proteins modulated by aging and eff^RNAi^ (Supplementary Table S4). The threshold of log_2_FC>0.3 and <-0.3 was further applied for selecting up- and down-regulated proteins.

**Fig. 7.**
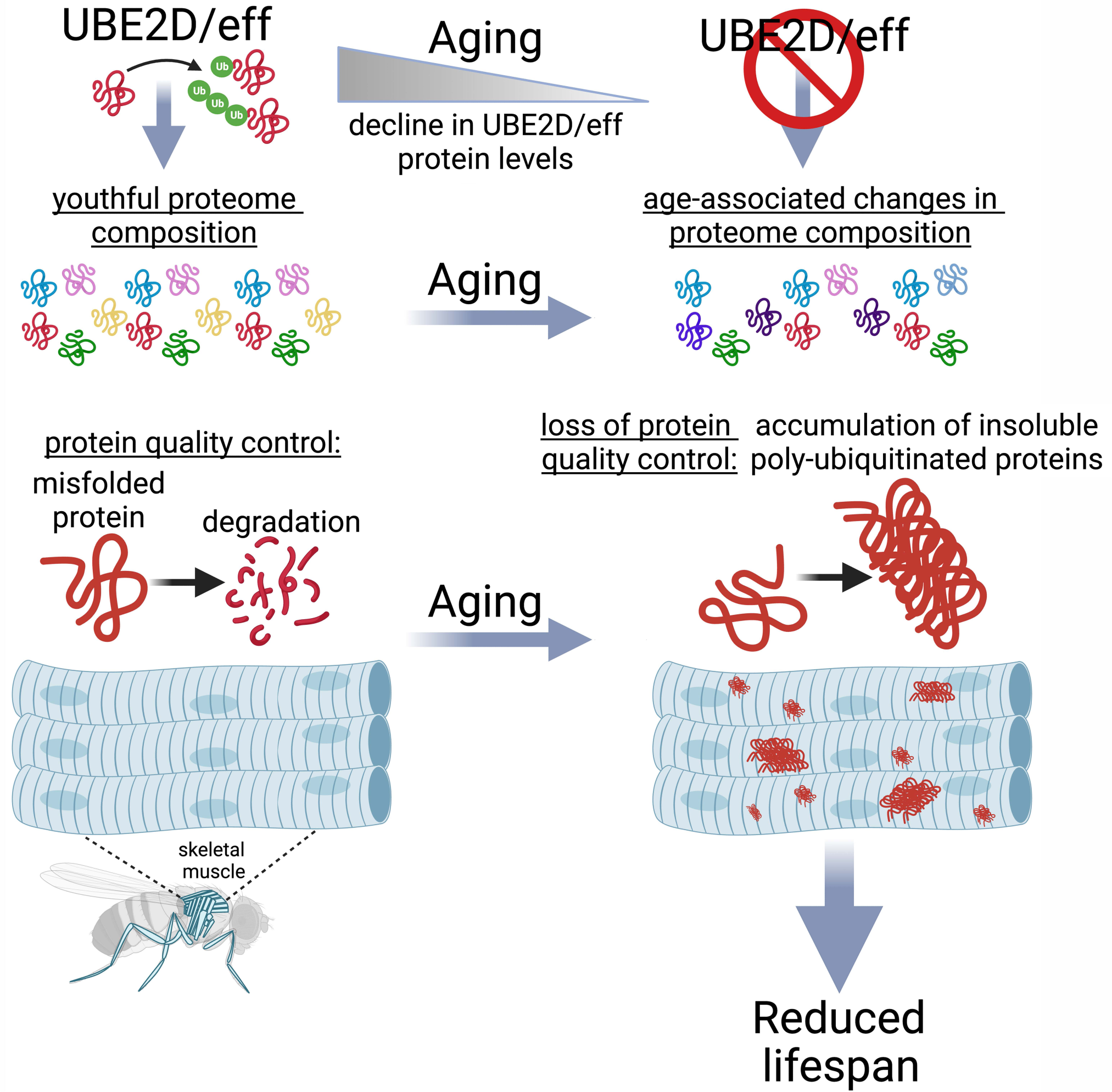
The E2 enzyme UBE2D/eff is necessary to preserve proteostasis and to maintain a youthful proteome composition in skeletal muscle during aging. The ubiquitin-conjugating enzyme UBE2D/eff has a key role in proteostasis in skeletal muscle: UBE2D/eff levels decline during aging and reproducing such UBE2D/eff knockdown from a young age causes a loss in protein quality control and a consequent precocious surge in the levels of insoluble poly-ubiquitinated proteins, which normally accumulate only in old age because of age-associated defects in proteostasis. Proteomics surveys indicate that UBE2D/eff knockdown rewires the proteome similar to aging, and that the UBE2D/eff^RNAi^-induced changes are rescued by transgenic expression of its human homolog UBE2D2. Altogether, these findings indicate that UBE2D/eff is necessary to maintain a youthful proteome and to ensure muscle protein quality control during aging.

Proteins that were upregulated by both eff RNAi and aging (the largest class, 514 proteins) were enriched for regulators of glutathione metabolism, chaperones, proteasome components, and vesicle transport (Fig. 6B) and included Arc1 and Arc2. Commonly downregulated proteins (186 proteins) included components of mitochondrial ribosomes and late endosomes, SNARE proteins, and proteases/peptidases (Fig. 6C). The discordantly regulated proteins were enriched for oxidoreductases, mitochondrial and endoplasmic reticulum components, and secreted proteins (106 proteins that are upregulated by aging but downregulated by eff RNAi; Fig. 6A), and regulators of cytoplasmic translation and mRNA splicing (196 proteins that are downregulated by aging but upregulated by eff RNAi; Fig. 6D).

Similar results were also obtained with Venn diagrams (Fig. 6E), which identified the proteins with overlapping, consistent, or discordant regulation by aging versus eff^RNAi^. The threshold of log_2_FC >0.3 and <-0.3 was used as a criterion for the selection of significantly up- and down-regulated proteins for these analyses (Fig. 6E). In this case, 23.9% of significantly regulated proteins (141) were upregulated by both aging and eff^RNAi^, 7.3% (43) were consistently downregulated by both, and 3.2% (19) were oppositely regulated (Fig. 6E).

Altogether, these analyses indicate that UBE2D/eff^RNAi^ causes proteomic changes that recapitulate in part those induced by aging, indicating that UBE2D/eff is necessary to maintain a youthful proteome composition.

## DISCUSSION

In this study, we have examined the role of the 21 *Drosophila* E2 ubiquitin-conjugating enzymes in modulating polyglutamine protein aggregation. By analyzing pathogenic huntingtin in the *Drosophila* retina, we found that E2s have diverse functions in regulating Htt-polyQ-GFP. Specifically, the knockdown of many E2s reduces polyglutamine protein aggregates (Fig. 1), suggesting that ubiquitination by these E2s promotes sequestration of pathogenic huntingtin into aggresomes, as found in *C. elegans* (*39*). Aggregation of Htt has been previously found to decrease its toxicity (*67, 68*), and therefore E2s that drive Htt aggregation may protect from polyglutamine disease via this mechanism. However, there were also RNAi interventions for other E2s, such as eff/UBE2D, and associated enzymes (DUBs and E3s) that increased the levels of high-molecular-weight assemblies of Htt-polyQ-GFP (Fig. 1-2), suggesting that these E2s and interacting partners are necessary for Htt-polyQ-GFP degradation. Likewise, eff/UBE2D knockdown compromises protein quality control also in *Drosophila* skeletal muscle (Fig. 3-4), as indicated by the accumulation of insoluble poly-ubiquitinated proteins, and this is in part re-established via overexpression of one of its human homologs, UBE2D2.

Considering that UBE2D/eff is one out of 21 different E2 ubiquitin-conjugating enzymes in Drosophila, it is unlikely that UBE2D/eff RNAi alone reduces the total ubiquitination capacity of the cell. In fact, UBE2D/eff knockdown paradoxically increases the levels of poly-ubiquitinated proteins, and such a generalized increase may result from an overall derangement in protein quality control. In agreement with this scenario, the protein categories that are modulated by UBE2D/eff^RNAi^ include regulators of proteostasis such as proteasome components and chaperones (Fig. 5). Therefore, derangement in the levels of a wide range of proteostasis regulators may lead to a generalized loss of protein quality control and to the accumulation of insoluble poly-ubiquitinated proteins upon UBE2D/eff RNAi.

Previously, UBE2D ubiquitin-conjugating enzymes (UBE2D1/2/3/4) were found to contribute to several cellular processes that impact proteostasis. These include the association of UBE2Ds with the E3 ubiquitin ligase CHIP to promote the degradation of misfolded proteins (*27*). Likewise, UBE2Ds were found to confer resistance to elevated temperatures in yeast by promoting the selective ubiquitination and degradation of misfolded proteins (*31, 69*). Moreover, UBE2Ds can interact with the E3 ligase parkin to promote mitophagy (*29*) and with the E3 ligase RNF138 to mediate DNA repair (*70*). In addition to these cellular functions, UBE2D also plays a key role in protein import into peroxisomes (*35, 71-74*).

We have found that eff/UBE2D knockdown leads to the accumulation of poly-ubiquitinated proteins and that eff/UBE2D is an E2 ubiquitin-conjugating enzyme that is key for maintaining protein quality control during aging. Because eff/UBE2D knockdown impedes eff/UBE2D-mediated ubiquitination, the poly-ubiquitinated proteins that accumulate upon eff/UBE2D RNAi likely consist of proteins different from eff/UBE2D ubiquitination substrates. How eff/UBE2D knockdown deranges proteostasis and promotes the accumulation of poly-ubiquitinated proteins may occur via multiple mechanisms. One possibility is that proteostasis is deranged by eff/UBE2D knockdown because of the previously-reported roles of eff/UBE2D in mitophagy (*29*) and in the removal of misfolded proteins (*31, 69*) in concert with the E3 ubiquitin ligase CHIP (*27*). Accumulation of such faulty proteins may clog or burden the proteasome and hence generally impair the degradation of poly-ubiquitinated proteins.

In an alternative model, proteostasis and survival are reduced by eff/UBE2D knockdown because of the accumulation of specific UBE2D protein targets that directly perturb these processes. While many substrates of eff/UBE2D ubiquitination accumulate and may play such role, Arc1 and the related Arc2 protein are of particular interest. Specifically, Arc1 and Arc2 protein levels increase upon UBE2D/eff knockdown and, conversely, they are reduced by hUBE2D2. Interestingly, Arc1 upregulation was found to be cytotoxic and to drive neurodegeneration in a *Drosophila* Alzheimer’s disease model (*64*). This suggests that the accumulation of Arc1/2 in response to eff/UBE2D knockdown may derange proteostasis and reduce survival. Arc1 and Arc2 protein levels increase also in response to aging in skeletal muscle (Fig. 6). More globally, eff RNAi induces many proteomic changes reminiscent of aging: ∼70% of the proteins that are significantly modulated by eff RNAi in young age are also consistently regulated by aging (Fig. 6). These findings, therefore, indicate that UBE2D/eff maintains a youthful proteome and that the age-dependent decline in UBE2D/eff protein levels (Fig. 3) may contribute to the proteomic changes that occur with aging.

In addition to reshaping the composition of the muscle proteome and impairing proteostasis, we have found that muscle-specific eff/UBE2D knockdown reduces organismal survival: while muscle-specific eff/UBE2D RNAi reduces lifespan, this is partially rescued by concomitant expression of an RNAi-resistant human UBE2D2 transgene which re-establishes the physiological levels of eff/UBE2D-regulated targets such as Arc1/2. These findings, therefore, reinforce the notion that skeletal muscle is a key tissue that influences systemic aging, and that preserving muscle protein quality control is necessary for optimal organismal survival (*25, 47, 57, 75-78*).

In summary, our study highlights the important role of the ubiquitin-conjugating enzyme eff/UBE2D in maintaining protein quality control during aging, suggesting that interventions that promote UBE2D function may contrast age-related diseases that arise from the loss of proteostasis.

## Acknowledgments

We thank the Light Microscopy facility at St. Jude Children’s Research Hospital. *Drosophila* stocks were provided by the VDRC, NIG-Fly, and the Bloomington stock centers. The scheme was drawn with BioRender.

## Funding

Work in the Demontis lab is supported by the National Institute on Aging of the NIH (R01AG055532 and R21AG079267) and the Alzheimer’s Association (AARG-NTF-22-973220). The Peng lab is supported by the NIH (RF1AG068581). The content is solely the responsibility of the authors and does not necessarily represent the official views of the National Institutes of Health. Research at St. Jude Children’s Research Hospital is supported by the ALSAC.

## Author contributions

L. C. H. did most of the experiments and data analysis, with help from K.N., J.J., and A.S.; K.K. and V.P did the TMT mass spectrometry and corresponding data analyses; J.P. supervised the mass spectrometry studies; F.D. supervised the project and wrote the manuscript.

## Competing interests

The authors declare that they have no competing interests.

## Data and materials availability

All data are available within the main text, figures, supplementary tables 1-4, and the source data file. The TMT mass spectrometry proteomics data have been deposited to the ProteomeXchange Consortium via the PRIDE partner repository and are accessible with the dataset identifiers PXD042345 and PXD045713.

## MATERIALS AND METHODS

### *Drosophila* husbandry and stocks

Flies were kept (∼25 flies/tube) at 25°C, 60% humidity, and a 12h/12h light-dark cycle in tubes containing cornmeal/soy flour/yeast fly food. Survival analysis was done at 25°C and the fly food was changed every 2-3 days (*79*). All experiments were done with male flies. Fly stocks were obtained from the Bloomington Drosophila Stock Center (BDSC), the Vienna Drosophila Resource Center (VDRC), and the National Institute of Genetics (NIG-Fly, Japan). The following fly stocks were utilized: *MhcF3-Gal4* ((*25, 80*) BL#38464), control *UAS-white^RNAi^*(BL#33623 and v30033), *UAS-luciferase^RNAi^* (BL#31603), *UAS-eff^RNAi^* (BL#35431, #7425R-2, and v26012), *UAS-mcherry* (BL#35787), *UAS-hUBE2D2* (BL#76819), and *UAS-mcherry^RNAi^* (BL#35785). *GMR-Gal4* and UAS-Htt-72Q-GFP flies were previously described (*41*). The fly stocks used for the screen in Fig. 1 are reported in Supplementary Table 1.

### Whole-mount immunostaining of *Drosophila* skeletal muscle

The immunostaining of flight skeletal muscle was done as previously described (*47, 81, 82*). In brief, thoraces were dissected, fixed for 30 minutes in PBS with 4% paraformaldehyde and 0.1% Triton X-100 at room temperature, washed >3 times in PBS with 0.1% Triton X-100 at room temperature, and immunostained overnight at 4°C with rabbit anti-poly-ubiquitin (FK2; Enzo Life Sciences #BML-PW8810-0100). After washes with PBS with 0.1% Triton X-100, the samples were incubated with secondary antibodies and Alexa635-phalloidin for 2 hours at room temperature, washed, and mounted in an antifade medium. Subsequently, the samples were imaged on a Nikon C2 confocal microscope. Image analysis was done with ImageJ.

### Analysis of pathogenic Huntingtin aggregation

Pathogenic huntingtin-polyQ72-GFP protein aggregates (*41*) were imaged with an epifluorescence ZEISS SteREO Discovery V12 microscope with consistent exposure time and settings. The acquired grayscale images were then analyzed in an automated manner by using Cell Profiler 3.0.0 (cellprofiler.org) to determine the number and/or total area of protein aggregates (Huntingtin-polyQ72-GFP speckles) normalized by the retinal tissue area. Background fluorescence not corresponding to Huntingtin-polyQ72-GFP aggregates was excluded by thresholding. This analysis was done with male flies after aging at 25°C for 30 days; the same conditions were applied to all the samples and the respective controls in any given experiment. The raw values for each set of flies were normalized by the corresponding raw values of the negative control RNAi intervention from the same fly stock collection.

### Western blots for detergent-soluble and -insoluble fractions

Western blots for detergent-soluble and insoluble fractions were done as before (*47, 56, 83*). In brief, thoraces were dissected from 20 male flies/sample and homogenized in ice-cold PBS with 1% Triton X-100 containing protease and phosphatase inhibitors. Homogenates were centrifuged at 14,000 rpm at 4°C and supernatants collected (Triton X-100 soluble fraction). The remaining pellet was washed in ice-cold PBS with 1% Triton X-100. The pellet was then resuspended in RIPA buffer containing 8M urea and 5% SDS, centrifuged at 14,000 rpm at 4°C. The supernatants (Triton X-100 insoluble fraction) were collected and analyzed on 4-20% SDS-PAGE with anti-ubiquitin (Cell Signaling Technologies P4D1, #3936) and anti-Ref(2)P/p62 (Abcam #178840) antibodies.

### Western blots

For the western blot analysis of flies that express Hungtingtin-polyQ72-GFP, 30-day-old flies (n=5/sample) were homogenized in RIPA buffer containing 8M urea and 5% SDS, and centrifuged at 14,000 rpm for 10 minutes at 4°C. The supernatants were collected and analyzed on 4-20% SDS-PAGE with anti-GFP antibodies (Cell Signaling Technologies D5.1, #2956). Anti-α-tubulin antibodies (Cell Signaling Technologies, #2125) were used as loading controls.

### qRT-PCR

qRT-PCR was performed as previously described (*26, 84-88*). Total RNA was extracted with the TRIzol reagent (Life Technologies) from *Drosophila* thoraces, consisting primarily of skeletal muscle, from >20 male flies/replicate, followed by reverse transcription with the iScript cDNA synthesis kit (Bio-Rad). qRT-PCR was performed with SYBR Green and a CFX96 apparatus (Bio-Rad). Three biological replicates were used for each genotype and time point. Alpha-tubulin at 84B (*Tub84B*) was used as a normalization reference. Whole flies at 1 week of age were utilized for the qRT-PCR in Fig.1D to detect *Htt* and *GFP* levels. The comparative C_T_ method was used for the relative quantitation of mRNA levels. The following qRT-PCR oligos were used:

*eff*: 5’-CAATAATGGGCCCGCCGGA-3’ and 5’-GCGCGTTGTAAAAGCCACTT-3’

*Tub84B*: 5’-GTTTGTCAAGCCTCATAGCCG-3’ and 5’-GGAAGTGTTTCACACGCGAC-3’

*GFP*: 5’-ACGTAAACGGCCACAAGTTC-3’ and 5’-CTTCATGTGGTCGGGGTAGC-3’

*Htt (exon 1)*: 5’-CCTGGATCCCTGGTGAGCAA-3’ and 5’-GCTGAACTTGTGGCCGTTTA-3’

### Protein sample preparation, protein digestion, and peptide isobaric labeling by tandem mass tags

For each TMT *Drosophila* sample, 50 thoraces (consisting primarily of skeletal muscle) from 1-week-old and 8-weeks-old male *w^1118^* flies (Fig. 3) and from 2-weeks-old male flies of the genotypes indicated (Fig. 5) were collected and homogenized in 8M urea lysis buffer (50 mM HEPES, pH 8.5, 8 M urea) (*89*). After homogenization with zirconium beads in a NextAdvance bullet blender, 0.5% sodium deoxycholate was added to the tissue homogenates, which were then pelleted to remove fly debris. The resulting supernatant was submitted for TMT mass spectrometry and digested with LysC (Wako) at an enzyme-to-substrate ratio of 1:100 (w/w) for 2 hours in the presence of 1 mM DTT. Following this, the samples were diluted to a final 2 M Urea concentration with 50 mM HEPES (pH 8.5), and further digested with trypsin (Promega) at an enzyme-to-substrate ratio of 1:50 (w/w) for at least 3 hours. The peptides were reduced by adding 1 mM DTT for 30 min at room temperature (RT) followed by alkylation with 10 mM iodoacetamide (IAA) for 30 minutes in the dark at RT. The unreacted IAA was quenched with 30 mM DTT for 30 minutes. Finally, the digestion was terminated and acidified by adding trifluoroacetic acid (TFA) to 1%, desalted using C18 cartridges (Harvard Apparatus), and dried by speed vac. The purified peptides were resuspended in 50 mM HEPES (pH 8.5) and labeled with 16-plex Tandem Mass Tag (TMT) reagents (ThermoScientific) following the manufacturer’s recommendations and our optimized protocol (*55*).

### Two-dimensional HPLC and mass spectrometry

The TMT-labeled samples were mixed equally, desalted, and fractionated on an offline HPLC (Agilent 1220) by using basic pH reverse-phase liquid chromatography (pH 8.0, XBridge C18 column, 4.6 mm × 25 cm, 3.5 μm particle size, Waters). The fractions were dried and resuspended in 5% formic acid and analyzed by acidic pH reverse phase LC-MS/MS analysis. The peptide samples were loaded on a nanoscale capillary reverse phase C18 column (New objective, 75 um ID × ∼25 cm, 1.9 μm C18 resin from Dr. Maisch GmbH) by an HPLC system (Thermo Ultimate 3000) and eluted by a 60-min gradient. The eluted peptides were ionized by electrospray ionization and detected by an inline Orbitrap Fusion mass spectrometer (ThermoScientific). The mass spectrometer was operated in data-dependent mode with a survey scan in Orbitrap (60,000 resolution,

- × 10^6^ AGC target and 50 ms maximal ion time) and MS/MS high-resolution scans (60,000 resolution,
- × 10^5^ AGC target, 120 ms maximal ion time, 32 HCD normalized collision energy, 1 *m*/*z* isolation window, and 15 s dynamic exclusion).

### MS data analysis

The MS/MS raw files were processed by the tag-based hybrid search engine, JUMP (*90*). The raw data were searched against the UniProt human and *Drosophila* databases concatenated with a reversed decoy database for evaluating false discovery rates. Searches were performed by using a 15-ppm mass tolerance for both precursor and product ions, fully tryptic restriction with two maximal missed cleavages, three maximal modification sites, and the assignment of *a*, *b*, and *y* ions. TMT tags on Lys and N-termini (+304.20715 Da) were used for static modifications and Met oxidation (+15.99492 Da) was considered as a dynamic modification. Matched MS/MS spectra were filtered by mass accuracy and matching scores to reduce protein false discovery rate to ∼1%. Proteins were quantified by summing reporter ion intensities across all matched PSMs using the JUMP software suite (*91*). Categories enriched in protein sets were identified with DAVID (*92*).

### Quantification and statistical analyses

All experiments were performed with biological triplicates unless otherwise indicated. The unpaired two-tailed Student’s *t*-test was used to compare the means of two independent groups to each other. One-way ANOVA with Tukey’s post hoc test was used for multiple comparisons of more than two groups of normally distributed data. Survival data was analyzed with OASIS 2 (*93*) by using log-rank tests. The “*n*” for each experiment can be found in the figures and/or figure legends and represents independently generated samples, including individual flies for lifespan assays, and batches of flies or fly thoraces for other assays. Bar graphs represent the mean ±SEM or ±SD as indicated in the figure legend. Throughout the figures, asterisks indicate a significant *p*-value (\**p*<0.05). Statistical analyses were done with Excel and GraphPad Prism.

### Description of the Supplementary Data

**Supplementary Table 1. RNAi screen for E2 ubiquitin-conjugating enzymes and associated E3 ubiquitin ligases that regulate pathogenic huntingtin-polyQ72-GFP protein aggregates in Drosophila.** Each E2/E3 was analyzed with multiple RNAi lines, and n=5 biological replicates were examined for each RNAi line.

**Supplementary Table 2. Mass spectrometry analysis of age-induced protein changes in Drosophila skeletal muscle.** Tandem mass tag (TMT) mass spectrometry identifies age-related protein changes in the skeletal muscle (thoraces) of old vs. young *w^1118^* male flies (8 vs. 1 week old). A tab also reports the list of age-regulated deubiquitinating enzymes (DUBs), E1s, E2s, and E3 enzymes.

**Supplementary Table 3. Mass spectrometry analysis of protein changes induced by eff/UBE2D knockdown in Drosophila skeletal muscle, and their rescue by co-expression of human UBE2D2.** Tandem mass tag (TMT) mass spectrometry identifies protein changes induced in skeletal muscle by eff RNAi (compared to control interventions) and their rescue by co-expression of hUBE2D2 (the human homolog of eff).

**Supplementary Table 4. Cross-comparison of proteomic changes induced by aging vs. eff RNAi in skeletal muscle.** Cross-comparison of the mass spectrometry data from Supplementary Tables 2 and 3.

**Supplementary Table 5. Source data and full scans of western blots.**

## Notes

### Competing Interest Statement

The authors have declared no competing interest.

### Summary of Updates

The results and method sections were updated for clarity; Figure 6 was revised by adding panel Fig. 6E.

